# Differentiating hallucination proneness dimensions through resting state alpha dynamics

**DOI:** 10.1101/2024.11.06.622288

**Authors:** Hanna Honcamp, Philip O’Donnell, Ana P. Pinheiro, Nelson J. Trujillo-Barreto, Michael Schwartze, Wael El-Deredy, Sonja A. Kotz

## Abstract

**Background:** Hallucination-like experiences (HLEs) are untriggered sensory perceptions linked to externalizing bias - the misattribution of self-generated sensory experiences to an external source. The vulnerability to HLEs, i.e., hallucination proneness (HP), is typically assessed by the Launay-Slade Hallucination Scale (LSHS). A recent LSHS factor analysis revealed four distinct HP dimensions: Multisensory HLEs, Auditory daydreaming, Vivid thoughts and inner speech, and Personified HLEs. The current study assesses whether these HP dimensions map onto distinct patterns of resting state brain dynamics in the alpha frequency band due to its modulatory role in attention and perception.

**Methods:** We used a Hidden semi-Markov Model to segment continuous RS alpha activity into nine recurrent brain states and extracted the total number of transitions (TT) and the number of visits per state (SV). We assessed how the HP dimensions relate to these metrics, calculated across the entire 3-minute recording and within shorter sliding windows to capture finer temporal changes.

**Results:** All HP dimensions and increased RS time correlated with increased TT. Increased Personified HLEs scores linked to different time-dependent changes of SV in two states (SV to state 5 decreased over time, while visits to state 9 increased), highlighting distinct alpha dynamics in high- and low-hallucination prone individuals.

**Conclusions:** Increased TT could indicate frequent attentional switches between internal and external states. Different SV patterns related to higher Personified HLEs scores suggest unstable source monitoring, potentially inducing an externalizing bias. These findings provide novel predictors of HP dimensions, revealing distinct neural profiles associated with different vulnerability profiles.

## Introduction

Hallucination-like experiences (HLEs) are transitory, untriggered sensory percepts that show in the general population. Increased HLE severity and frequency, i.e., hallucination proneness (HP), can indicate a heightened risk of developing clinically relevant hallucinations (Johns & Van Os, 2001; Larøi et al., 2019; Yates et al., 2021). The Launay Slade Hallucination Scale (LSHS) is a widely used measure of HP (Launay & Slade, 1981). While some LSHS items pertain to HP in specific modalities (e.g., auditory, visual), previously explored LSHS factorial structures show that HP is multidimensional and multisensory, suggesting that HP may not be adequately represented by modality-specific composite scores (Fonseca-Pedrero et al., 2010; Honcamp, Goller, et al., 2024; Vellante et al., 2012). This implies that different HP dimensions may have distinct neurophysiological underpinnings.

HLEs and HP correlate with aberrant functional connectivity in the default-mode network (DMN) and altered interaction between the DMN and other resting state networks (RSNs) (Kottaram et al., 2019; van Lutterveld et al., 2014; Weber et al., 2020). Given the fluctuating and untriggered nature of HLEs, analyses of temporal dynamics of spontaneous, task-free (resting state, RS) brain activity could uncover neurophysiological correlates of HP (Alderson-Day, McCarthy-Jones, & Fernyhough, 2015; Geng et al., 2020; Honcamp et al., 2022; Northoff & Qin, 2011).

Recent research used high-temporal resolution methods like magneto-/electroencephalography (M/EEG) and computational modelling to capture rapid neural fluctuations of the resting brain (Baker et al., 2014; Vidaurre et al., 2016; Woolrich et al., 2013). The generative Hidden semi-Markov Model (HsMM) segments RS brain activity into a sequence of distinct, recurrent states, considering realistic assumptions about network switching and long-range dependencies of neural time series data (Honcamp et al., 2022; Trujillo-Barreto et al., 2024). Each state is characterized by a specific pattern of RSN functional connectivity, reflecting the brain’s baseline temporal dynamics (Baker et al., 2014). The temporal characteristics of HsMM states, such as their transition frequency and number of visits, can be linked to cognition and behavior as well as to vulnerability to various neuropsychiatric conditions (Honcamp, Duggirala, et al., 2024; Lin et al., 2022; Ou et al., 2015; Puttaert et al., 2020; Zhang et al., 2022). A higher transition frequency was observed during hallucinations compared to rest, which was interpreted as more erratic brain activity during hallucinations (Marschall et al., 2023). Moreover, schizophrenia has been associated with less frequent transitions in and out of specific networks including the DMN, but once visited, the network activation duration was significantly increased (Kottaram et al., 2019).

Specific EEG frequency bands also capture important individual differences in cognition and behavior, and inform about increased HP (Ford et al., 2012; Honcamp, Duggirala, et al., 2024). Alpha band frequency fluctuations were linked with self-reflective thinking, sensory sensitivity, and attentional switching from external to internal states (Benedek et al., 2014; Hillebrand et al., 2012; Iemi et al., 2017; Webster & Ro, 2020). These processes may promote the misattribution of internally generated sensations to an external source, a potential key mechanism underlying HP (Badcock & Hugdahl, 2012; Brookwell, Bentall, & Varese, 2013). Alpha activity also tracks internally directed cognitive states, such as spontaneous thoughts and mind wandering, which share underlying neural correlates with HP (Fazekas, 2021; Groot et al., 2021; Zanesco, Denkova, & Jha, 2021). Time spent in the RS also correlates with internally directed attention as external constraints decrease over time (Christoff et al., 2016). Thus, alpha band dynamics and their change over time could indicate individual differences in HP.

The current study builds on and extends a novel LSHS factor structure to investigate the electrophysiological correlates of four HP dimensions: Multisensory HLEs; Auditory daydreaming, Vivid thoughts and inner speech, and Personified HLEs (Honcamp, Goller, et al., 2024). We focused on HsMM-inferred transition frequency and state visits in the alpha frequency band and examined their change over time to account for spontaneous fluctuations in functional connectivity and cognition.

## Materials and methods

### Participants and procedure

The sample included data of 65 individuals from the general population (50 females, 15 males, mean age = 21.57, SD = 6.59, age range = 18 – 48). Participants provided written informed consent before participation. Ethical approval was granted by the Deontological Committee of the Faculty of Psychology at the University of Lisbon.

### Data aquisition and processing

Data were collected at the University of Lisbon, Portugal. The Portuguese version of the LSHS was used to measure HP (Castiajo & Pinheiro, 2017). The factor scores of each factor Multisensory HLEs (MS-HLEs), Auditory daydreaming (AD), Vivid thoughts and inner speech (VT-IS), and Personified HLEs (P-HLEs) were taken from an exploratory factor analysis of the LSHS (Honcamp, Goller, et al., 2024) (Supplementary Material A).

RS EEG data were recorded for three minutes in an eyes-closed condition using a 64-channel BioSemi Active system (BioSemi, Amsterdam, Netherlands) in an acoustically and electrically shielded booth. Participants were instructed to minimize movements during data acquisition. EEG data were preprocessed in EEGLAB v2023. The preprocessing was adapted from the pipeline outlined in (Honcamp, Duggirala, et al., 2024) and included downsampling to 512 Hz, band-pass filtering between 1-40 Hz, interpolation of noisy channels, average re-referencing, Automatic Subspace Reconstruction, and Independent/Principal Component Analysis (ICA/PCA) to remove non-brain related components from the data. On average, 5.17 (SD = 3.13) noisy channels were interpolated, and 5.92 (SD = 2.00) components were removed per participant.

### HsMM analysis

Data preparation for HsMM modeling followed the data transformation steps in Honcamp et al., 2024b, including filtering to the frequency band of interest (alpha, 8-12 Hz), signal amplitude envelope extraction using the Hilbert transform method, signal normalization, logarithmic transformation, PCA, and downsampling. Twenty-five principal components were retained, which explained 90.64 % of the variance.

Currently, there is no consensus regarding the optimal number of states to be modeled in the HsMM (Honcamp, Duggirala, et al., 2024; Kottaram et al., 2019; Kotz et al., 2023; Vidaurre et al., 2016). However, it can be inferred by maximizing negative Free Energy (nFE), a model quality measure that balances the complexity and accuracy of the model (Trujillo-Barreto et al., 2024). Here, we trained separate HsMMs with 1 to 12 states for 15 randomly selected participants. Most models converged between 7 and 11 states (Supplementary material B). Thus, we chose 9 states as a compromise between participants. Final HsMM modeling of 9 states was performed on the concatenated and downsampled principal components of all 65 participants.

**Figure 1.**
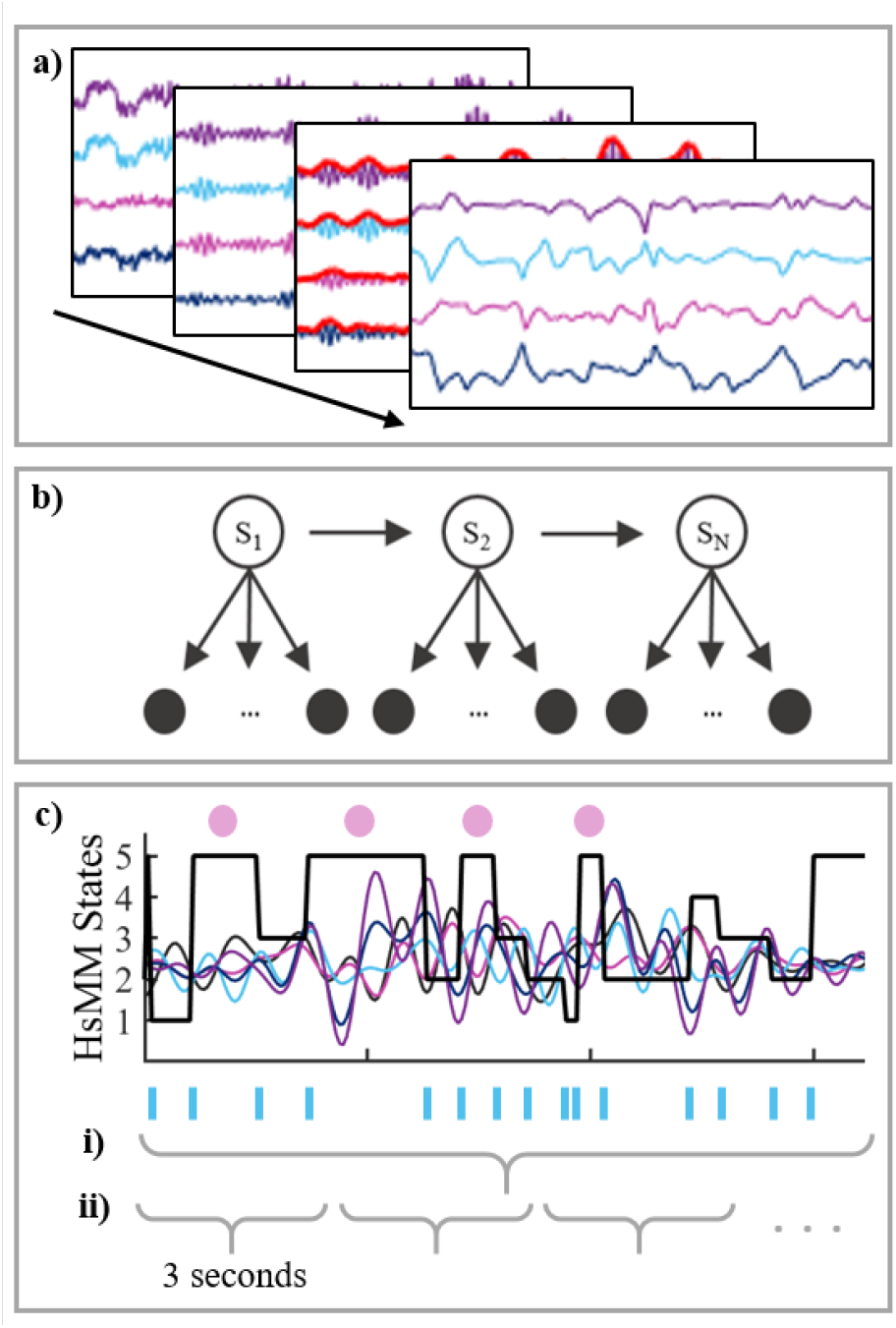
Data processing and analysis pipeline. **a) Resting state EEG data pre-processing.** The concatenated time series for all participants is filtered to the alpha band (8-12 Hz). The amplitude envelope is extracted using the Hilbert transform method, followed by normalization. The normalized envelopes are then log-transformed (not shown here), subjected to PCA (90.64% explained variance retained with 25 components), and downsampled to 64 Hz. **b) HsMM**. The concatenated data of all participants is modeled using a multivariate normal (MVN) emission model and a lognormal duration model. Nine states were identified. **c) Calculation of state metrics**. Temporal dynamics of the HsMM-inferred states were derived from each participant’s state sequence for **i)** the entire three-minutes recording and **ii)** a sliding window of three seconds. The blue lines represent the transitions between states, whereas the pink dots represent the state visits to an example state.

### Calculation of state metrics

The state sequence of each participant was used to calculate the total number of state transitions (TT), i.e., the number of times a participant transitions across all 9 states and the number of state visits (SV), i.e., the number of times a participant visits a particular state. Additionally, we used a sliding window approach to quantify the dynamic change of TT and SV over time using a window size of three seconds with no overlap. Thus, TT and SV were obtained i) for the whole three-minute recording period, and ii) for different time windows allowing the quantification of change across time.

### Statistical analysis

Statistical analyses were performed in IBM SPSS v26 (IBM Corp., 2019) and Matlab R2020b (MathWorks Inc.). To assess the relationship between the states’ dynamic features and the HP dimensions, we ran several simple linear regression models with TT or SV as the dependent variable and one of the HP dimensions (MS-HLEs, AD, VT-IS, or P-HLEs) as predictor. To assess the dynamic change of TT and SV over time and whether this change depends on an individual’s score on a particular HP factor, we first performed a median split for each HP factor to assign individuals into a low or high scoring group. Subsequently, several multiple linear regression models were performed with either TT or SV as dependent variable, and time, group, and the interaction between time and group as predictors.

## Results

The HsMM-inferred maps of all 9 states are depicted in Supplementary Figure 3.

### Total transitions and state visits

The linear regression models with TT as dependent variable indicated that the total number of transitions within the whole three-minute recording period significantly increased as the scores on all four factors, MS-HLEs, AD, VT-IS, and P-HLEs increased (Figure 3, Table 1). To account for multiple hypothesis tests, the Benjamini-Hochberg (BH) procedure was applied to control the false discovery rate (FDR, Benjamini & Hochberg, 1995). After FDR correction, the linear relationship between all factors and TT remained significant (MS-HLE: R^2^ = .089, t= 2.488, adj. p = .032; AD: R^2^ = .099, t = 2.635, adj. p = .044; VT-IS: R^2^ = .089, t = 2.485, adj. p = .021; P-HLE: R^2^ = .064, t = 2.077, adj. p = .042).

**Figure 2.**
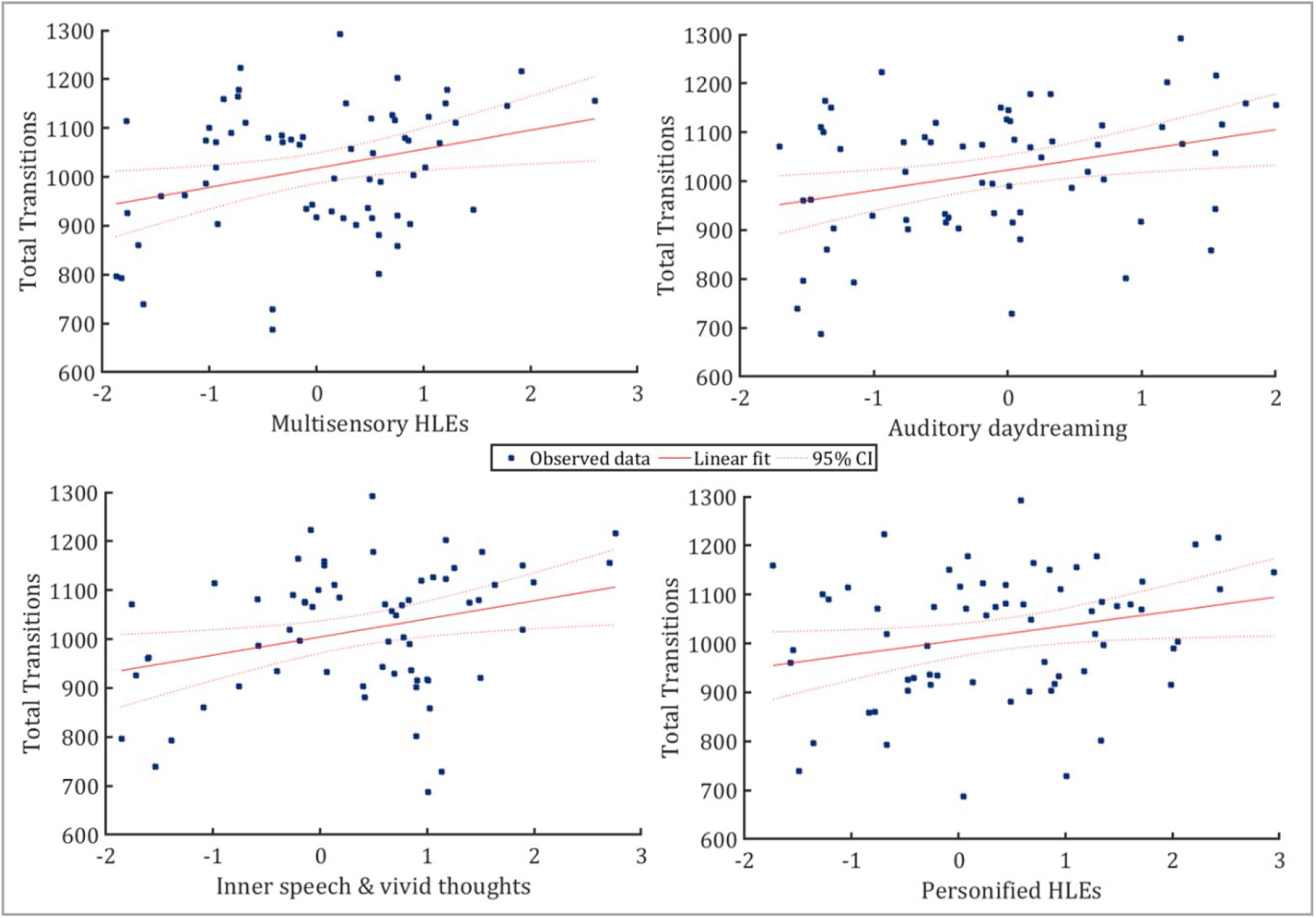
Linear relationship between the total transitions and each HP factor. Blue dots represent the total transitions across all states within the entire three-minutes recording for each participant; Bold red lines represent the fitted regression line; Dotted red lines represent the 95% confidence (CI) interval. Table 1 reports the corresponding statistics.

**Figure 3.**
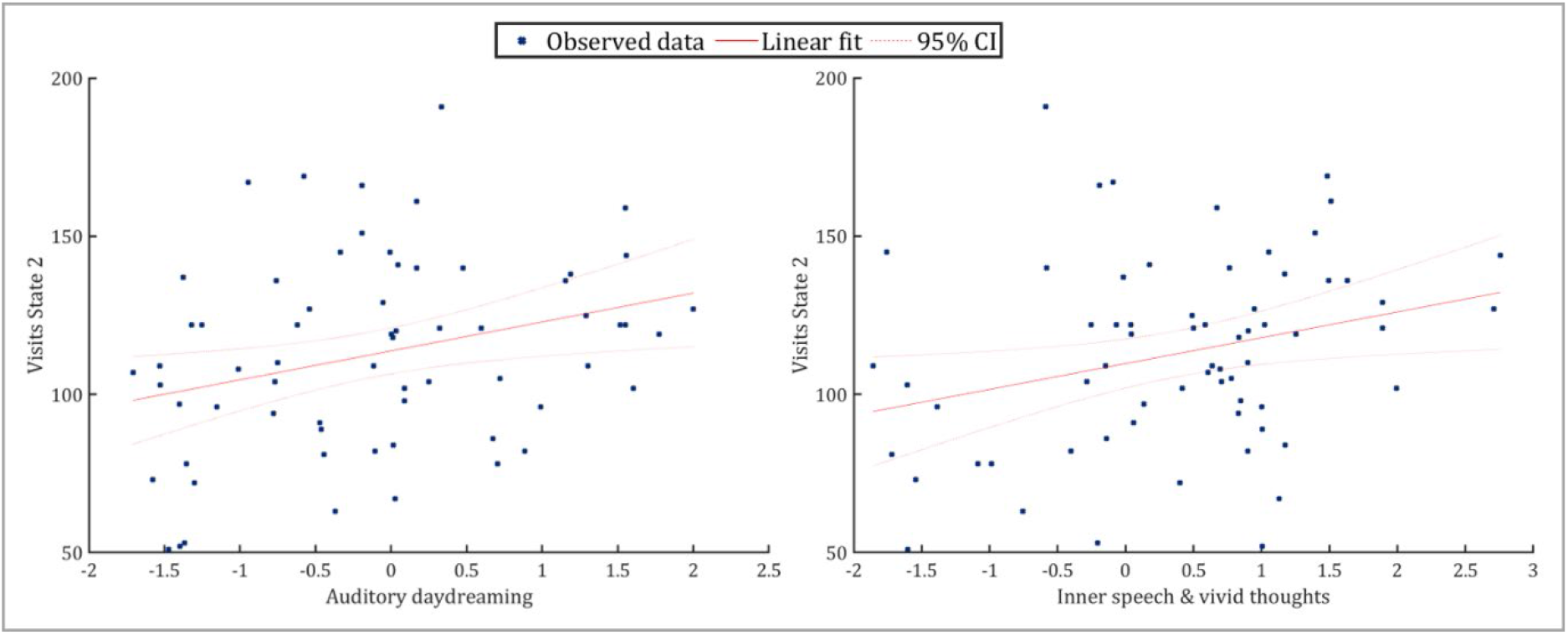
Linear relationship between the visits to state 2 and factors Auditory daydreaming and Vivid thoughts and inner speech. Blue dots represent the number of visits to state 2 within the entire three-minute recording for each participant; Bold red lines represent the fitted regression line; dotted red lines represent the 95% confidence (CI) interval. Table 2 reports the corresponding statistics.

**Table 1.** Results of the linear regression models: Total transitions and HP factors.

| Factor | R<sup>2</sup> | F | B | SE | t | p | Adj. p |
| --- | --- | --- | --- | --- | --- | --- | --- |
| MS-HLEs | .089 | 6.190 | 38.964 | 15.660 | 2.488 | .016* | .032* |
| AD | .099 | 6.943 | 41.445 | 15.729 | 2.635 | .011* | .044* |
| VT-IS | .089 | 6.176 | 37.012 | 14.893 | 2.485 | .016* | .021* |
| P-HLEs | .064 | 4.316 | 29.730 | 14.311 | 2.077 | .042* | .042* |

MS-HLEs = Multisensory HLEs, AD = Auditory daydreaming, VT-IS = Vivid thoughts and inner speech, P-HLEs = Personified HLEs; \*p < .05; Adj. p = p-values adjusted with the Benjamini-Hochberg procedure.

**Table 2.** Results of the linear regression models: Visits to state 2 and factors AD and VT_IS.

| Factor | $R^2$ | F | B | SE | t | p | Adj. p |
| --- | --- | --- | --- | --- | --- | --- | --- |
| AD | .090 | 6.220 | 9.153 | 4.670 | 2.494 | .015* | .060 |
| VT_IS | .080 | 5.502 | 8.149 | 3.474 | 2.346 | .022* | .030* |

AD = Auditory daydreaming, VT-IS = Vivid thoughts and inner speech. \* $p < .05$ ; reported p-values are uncorrected. Adj. $p = p$ -values adjusted with the Benjamini-Hochberg procedure.

To assess whether SV to any of the 9 states was predicted by the HP factor scores, separate linear regressions models were conducted. The relationship between visits to state 2 was predicted by the scores on factors AD and VT-IS (Figure 4, Table 2), however, only the linear relationship between VT-IT and visits to state 2 remained significant after controlling for FDR (R^2^ = .080, t = 2.346, Adj. p = .030).

**Figure 4.**
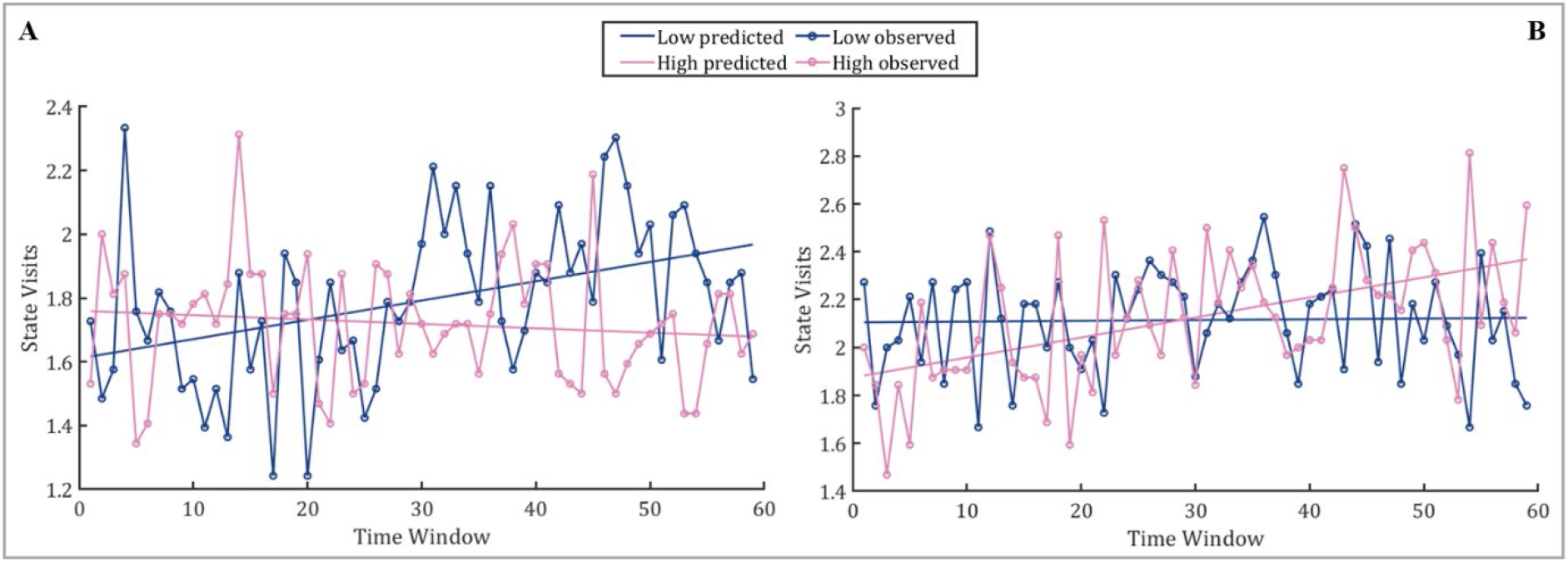
Group by time interaction – State visits and Personified HLEs. **A.** Visits to state 5; **B**. Visits to state 9. Solid lines represent the fitted linear regression line; Dotted lines represent the observed number of state transitions within each time window averaged for each group. Blue and pink colors depict the low-scoring and high-scoring groups, i.e., individuals scoring below or above the median for each factor.

### Change of total transitions and state visits over time

We also assessed whether the change of TT and SV over time, i.e., across sliding time windows of three seconds, differed between the low- and high-scoring group for each individual HP factor. For TT, none of the group-by-time interactions were significant (Supplementary Material C). The main effect of time was significant for factors MS-HLEs and AD, and approached significance for P-HLEs, indicating that TT tended to increase as a function of time. Moreover, the significant main effects of the group for factors MS-HLEs, AD, and P-HLEs support the positive association between TT and HP, which was stable across the 3-minute recording period.

The results further yielded a significant group by time interaction of SV to states 5 and 9 and P-HLEs, which remained significant after FDR correction (Figure 5, Table 3). While SV to state 5 decreased as a function of time for the high-scoring group, they increased for the low-scoring group (t = -2.890, adj. p = .017). We found an opposite pattern for state 9, where SV to state 9 increased over time for the high-scoring group, whereas the low-scoring group showed a decrease over time (t = 2.978, adj. p = .026). Notably, states 5 and 9 have distinct topographies with almost opposite activity distributions. While state 5 is characterized by higher positive voltages over temporal electrodes, state 9 shows higher positive voltages over frontal and posterior parietal electrodes.

**Table 3.** Results of the linear regression models: Visits to state 5 and 9 over time and Personified HLEs.

| Model summary |  |  | Coefficients |  |  |  |  |
| --- | --- | --- | --- | --- | --- | --- | --- |
| State | R<sup>2</sup> | F | B | SE | t | p | Adj. p |
| 5 | .004 | 4.879 |  |  |  |  |  |
| Intercept |  |  | 1.610 | .062 | 25.891 | 0 |  |
| Group |  |  | .150 | .089 | 1.687 | .092 |  |
| Time |  |  | .006 | .002 | 3.356 | < .001** |  |
| Group x Time |  |  | -.007 | .003 | -2.890 | < .005** | .017* |
| 9 | .005 | 6.330 |  |  |  |  |  |
| Intercept |  |  | 2.105 | .065 | 32.176 | 0 |  |
| Group |  |  | -.230 | .093 | -2.472 | .013* |  |
| Time |  |  | .000 | .002 | .170 | .865 |  |
| Group x Time |  |  | .008 | .003 | 2.978 | < .005** | .026* |

## Discussion

The current study investigated the relationship between temporal dynamics of RS alpha band activity and different dimensions of HP, namely M-HLEs, AD, VT-IS, and P-HLEs. To this end, an HsMM was applied to non-clinical participants’ concatenated data, which identified nine states with unique spatio-temporal dynamics. The analyses specifically addressed the relationship between each HP dimension and two HsMM-inferred state metrics: the total number of state transitions (TT) and the number of visits to a given state (SV) for the entire three-minute recording period, and over time, i.e., across subsequent time windows.

The results indicated that all four factors were associated with TT. This pattern persisted over time; both low- and high-scoring groups showed a similar TT increase across time windows. Additionally, VT-IS was associated with visits to state 2, while P-HLEs showed a differential pattern of visits to states 5 and 9 over time. While the low-scoring group’s SV to state 5 increased over time, the high-scoring group showed a decrease. The reverse pattern was observed for state 9, where the low-scoring group visited state 9 less as time passed, whereas the high-scoring group showed an increase in SV to state 9 across time.

### Attention to internal states, alpha, and externalization bias

During rest, external constraints on the brain (e.g., task and attentional demands) weaken, which facilitates spontaneous, stimulus- and task-unrelated thoughts, or mind wandering (Christoff et al., 2016). Stimulus- and task-unrelated thoughts are closely related to interoception. During rest, the interoceptive, internally directed attentional state, is periodically interrupted by an exteroceptive state, i.e., attention to the external environment (Fransson, 2005). Interestingly, individuals with psychosis showed an altered interaction between extero- and interoception, which could impair successful differentiation between internally and externally generated percepts (Damiani et al., 2022; Yao & Thakkar, 2022).

One of the most consistently reported electrophysiological correlates of interoception is a dynamic change of alpha power, especially over posterior parietal electrodes (Compton, Gearinger, & Wild, 2019; Dhindsa et al., 2019; Jin, Borst, & Van Vugt, 2019; Webster & Ro, 2020). Thus, high versus low alpha power in posterior regions could differentiate between internally and externally directed attentional states. Additionally, functional magnetic resonance imaging (fMRI) studies have consistently identified increased activation of the DMN, including the posterior and anterior cingulate cortices, as an indicator of spontaneous thoughts and interoception (Christoff et al., 2016; Knyazev et al., 2011; Kucyi & Davis, 2014; Mason et al., 2007).

Given the current focus on alpha-band dynamics, the increase in TT over time may reflect recurrent shifts in attention from exteroceptive to interoceptive states as external constraints diminish. This interpretation seems reasonable given that the data were recorded in a sound-shielded and visually static booth. HP was further associated with more frequent state transitions, suggesting that hallucination-prone individuals may experience more frequent shifts toward self-focused attention and mind-wandering (Fazekas, 2021; Perona-Garcelán et al., 2014).

Prior research supports the link between HP and more frequent state switching. A dynamic functional connectivity study by Marschall et al. (2023) using leading eigenvector dynamic analysis (LEiDA) with individuals reporting auditory-verbal hallucinations found that hallucination periods were characterized by more frequent state switching as compared to pure rest periods. More frequent state switching often coincided with decreased duration of brain states, i.e., the shorter their duration, the quicker the switch to another state. An association between HP and reduced state durations was found in fMRI-based dynamic functional connectivity studies. Weber et al. (2020) found that HP of schizophrenia patients was associated with reduced state durations in a state characterized by a strong anti-correlation between the DMN and task-positive networks. Geng et al. (2020) found a similar pattern, in which schizophrenia patients with hallucinations showed decreased duration of a state characterized by a DMN-language network anti-correlation as compared to patients without hallucinations. This suggests a consistent relationship between HP and rapid transitions between network connectivity patterns, aligning with the current findings.

The results regarding the time-dependent changes of SV are more complex. In an earlier study with a similar methodology but a different non-clinical sample, we reported an association between auditory(-verbal) HP and the duration of a state with topographical features similar to state 9 in the current study (Honcamp, Duggirala, et al., 2024). Source localization revealed that this state related to active sources in the posterior DMN, somatosensory, and auditory network hubs, in accordance with theories on the neurophysiological correlates of hallucinations and increasing HP (Alderson-Day, McCarthy-Jones, & Fernyhough, 2015; Honcamp, Duggirala, et al., 2024; Kottaram et al., 2019; van Lutterveld et al., 2014). Although the current study did not consider the states’ durations, the increased SV to state 9 over time for the high-hallucination prone group suggest this state’s relevance in HLEs vulnerability. Given that the factor P-HLEs includes two “auditory” items that were previously included in the auditory sum score used in (Honcamp, Duggirala, et al., 2024), the current results seem consistent with the earlier findings.

P-HLEs, which address the tendency to attribute sensory experiences to another person, is closely related to an externalizing bias, a cognitive mechanism shown in both clinical and non-clinical hallucinations (Brookwell, Bentall, & Varese, 2013). Evidence suggests that the bias to (falsely) attribute internally generated sensory stimuli to an external source could be a trait-like cognitive vulnerability that correlates with HP across a continuum (Brookwell, Bentall, & Varese, 2013; Livet et al., 2020; Woodward, Menon, & Whitman, 2007). The interaction between HP, externalizing bias, and self-focused attention becomes particularly interesting when considering salience, i.e., the importance or attention an individual assigns to a stimulus. In hallucination-prone individuals, internally generated sensory events, such as bodily sensations or thoughts, can be imbued with heightened salience, which amplifies externalizing bias (Kapur, 2003; Kowalski et al., 2021; Moseley, Fernyhough, & Ellison, 2013). However, it remains unclear whether the increased salience assigned to internally generated events precedes or follows a heightened internally directed attentional focus.

Assuming that the time spent in a task-free setting facilitates more frequent inward attentional shifts, state 9 could represent interoception. This interpretation would be consistent with the topography of state 9 (Supplementary Figure 3), showing high alpha-band activity over posterior parietal and frontal midline electrodes, and the source localization results reported in Honcamp, Duggirala, et al. (2024), indicating involvement of the DMN. The increased number of visits to state 9 as a function of time observed in the high-hallucination-prone group could thus indicate recurrent periods of internally directed attention, resulting in higher engagement of source monitoring in response to highly salient internally generated sensory events. This suggests a shift toward increased awareness of internal sensations and an increased need to determine whether they originate internally or externally (Ensum & Morrison, 2003; Kowalski et al., 2021). Conversely, state 5 could represent an inhibitory control or regulatory mechanism of salience attribution to internally or externally generated sensations. Decreased engagement of such processes, as indexed by the reduced number of visits to state 5 for the high-hallucination proneness group, could facilitate misattributing internally generated sensory experiences to an external source (Alderson-Day et al., 2019; Hugdahl, 2009; Lesh et al., 2011; Waters et al., 2012).

### Limitations and suggestions for future research

This study is the first to investigate the neurophysiological correlates of HP dimensions using the novel LSHS factor structure. Future research should validate these dimensions in different samples using Confirmatory Factor Analysis (Brown, 2015). Moreover, although research corroborated the relationship between HP, increased self-focus, altered source monitoring, and externalizing bias, some studies did not find supporting evidence for this (Alderson-Day et al., 2019; Ardizzi et al., 2016; Garrison et al., 2017). Additionally, the current study did not assess the participants’ propensity to engage in spontaneous thoughts, attention, and source monitoring directly. Future research should therefore integrate direct measures of attention and source monitoring with brain dynamics.

The current findings provide novel insight into the neurophysiological substrates of refined measure of non-clinical HP, especially regarding the tendency to attribute sensory experiences to another person (Honcamp, Goller, et al., 2024). However, the small effect sizes indicate that a large proportion of variance remains unexplained. Including potentially confounding variables (e.g., attentional performance, daydreaming propensity) could mitigate this problem in upcoming studies. Lastly, applying dynamic state-space models, such as the HsMM, to neuroimaging data with higher spatial resolution would strengthen the functional interpretation of identified frequency-dependent brain states.

## Conclusion

The current study sheds light on the neurophysiological correlates of non-clinical HP, focusing on temporal dynamics of EEG alpha-band activity and distinct LSHS factors: M-HLEs, AD, VT-IS, and P-HLEs (Honcamp, Goller, et al., 2024). The findings suggest that individuals scoring higher on any of the four factors exhibit more frequent state transitions, suggesting a greater tendency for internally focused attention and mind-wandering. Additionally, the association between specific patterns of state visits, particularly to states 5 and 9, and P-HLEs may reflect processes related to source monitoring and externalizing bias, which both have been implicated in hallucinatory vulnerability (Brookwell, Bentall, & Varese, 2013; Kowalski et al., 2021; Woodward, Menon, & Whitman, 2007). These results contribute to a more nuanced understanding of HP, guiding further research that includes direct measures of attention and source monitoring, and complementary neuroimaging methods to replicate and extend these findings.

## Ethics approval

Ethical approval was granted by the Deontological Committee of Faculty of Psychology at the University of Lisbon, Portugal.

## Consent statement

All participants provided written informed consent before participation.

## Data and code availability statement

The software used for HsMM implementation in this manuscript is available at https://github.com/daraya78/BSD. Anonymized data will be shared with other researchers upon reasonable request to the corresponding author.

## CRediT authorship contribution statement

All authors contributed to and have approved the final manuscript. HH: Conceptualization, Methodology, Software, Formal analysis, Data Curation, Writing - Original Draft, Writing – Review & Editing, Visualization; PO: Conceptualization, Methodology, Writing - Original Draft; APP: Resources, Writing – Review & Editing; MS: Writing – Review & Editing; WED: Writing – Review & Editing; NTB: Writing – Review & Editing; SAK: Conceptualization, Supervision, Resources, Writing – Review & Editing.

## Acknowledgements

We thank Maria Amorim for her contribution to the EEG data collection and Sabrina Benvenuti and Jan Ostermann for valuable data explorations during their research internships.

