## Supplementary Material for "Differentiating hallucination proneness dimensions through resting state alpha dynamics"

Dept. Neuropsychology and Psychopharmacology

Maastricht University

Universiteitssingel 40, 6229 ER Maastricht, the Netherlands

### A. Supplementary methods

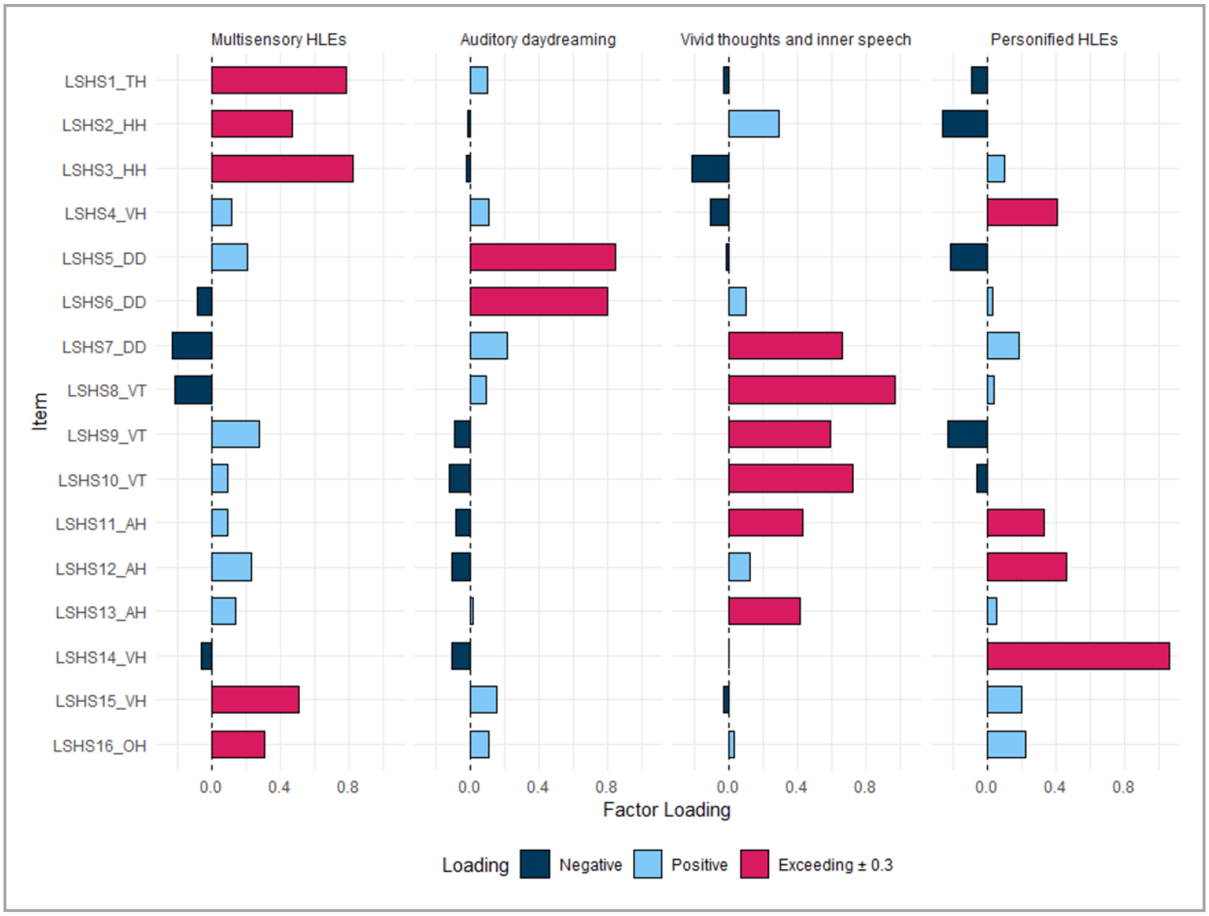


**Supplementary Figure 1.** Four-factor structure of the Launay-Slade Hallucination Scale. Factor loadings were taken from Honcamp et al. (2024). Dark and light blue bars indicate negative and positive loadings, respectively. Pink bars reflect loadings exceeding 0.3, referring to those items that were identified within each respective factor.

### B. Choosing the optimal number of states

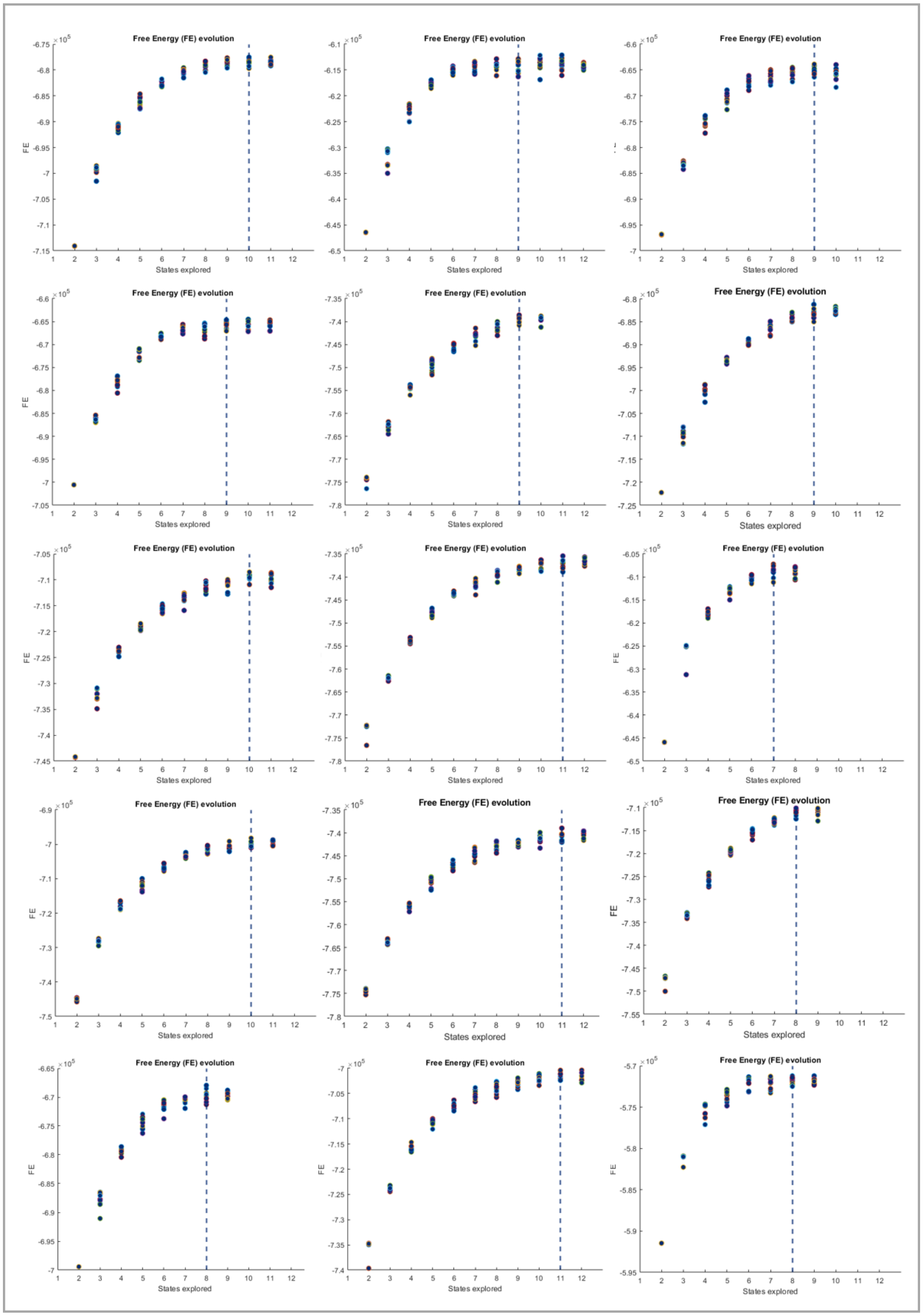

**Supplementary Figure 2.** Negative Free Energy evolution of a subset of participants. The individual graphs represent the negative Free Energy (nFE) evolution across different models with a different number of states (4-12). The dotted blue line indicates the optimal number of states. Individual data points show FE values for five model training iterations.

### C. Additional results

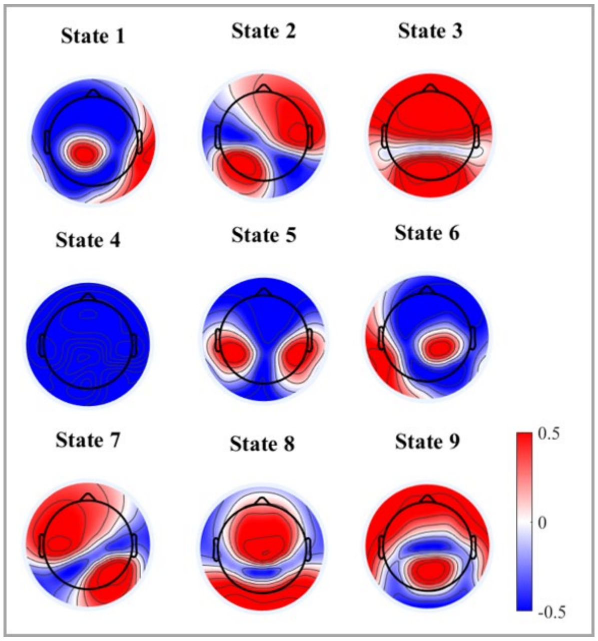


**Supplementary Figure 3.** State maps. The state maps depict the most representative topography for each HsMM-inferred state, obtained by multiplying the mean vector of the MVN distribution of each state with the PCA coefficients obtained during the data processing. The order of the states is arbitrary. The scale represents microvolts.

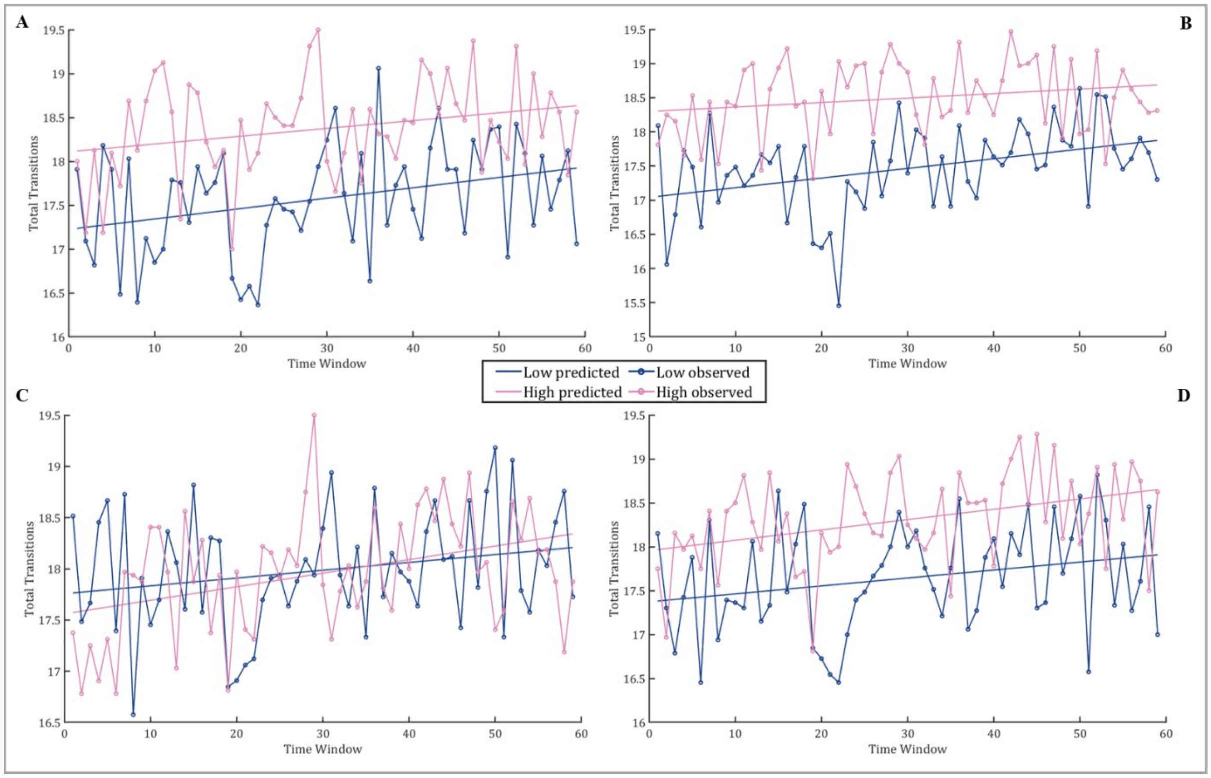


**Supplementary Figure 4.** Group by time interaction – Total transitions and HP factors. **A.** Multisensory HLEs. **B.** Auditory daydreaming. **C.** Vivid thoughts and inner speech. **D**. Personified HLEs. Solid lines represent the fitted linear regression line; Dotted lines represent the observed number of state transitions within each time window averaged for each group. Blue and pink colors depict the low-scoring and high-scoring groups, i.e., individuals scoring below or above the median for each factor.

**Supplementary Table 1.** Results of the linear regression models: Total transitions over time and HP factors

| | | Model summary | | Coefficients | | | | |
|---|---|---|---|---|---|---|---|---|
| **Factor** | | **$R^2$** | **F** | **B** | **SE** | **t** | **p** | **Adj. p** |
| MS-HLEs | | .013 | 16.628 | | | | | |
| | Intercept | | | 17.225 | .175 | 98.149 | .000 | |
| | Group | | | .884 | .250 | 3.533 | .000* | |
| | Time | | | .012 | .005 | 2.329 | .020* | |
| | Group x Time | | | -.003 | .007 | -.407 | .684 | .0912 |
| AD | | .020 | 26.506 | | | | | |
| | Intercept | | | 17.042 | .175 | 97.475 | .000 | |
| | Group | | | 1.256 | .249 | 5.040 | .000* | |
| | Time | | | .014 | .005 | 2.782 | .005* | |
| | Group x Time | | | -.008 | .007 | -1.043 | .297 | 1.188 |
| VT-IS | | .002 | 2.928 | | | | | |
| | Intercept | | | 17.757 | .176 | 100.644 | .000 | |
| | Group | | | -.197 | .251 | -.784 | .433 | |
| | Time | | | .008 | .005 | 1.492 | .136 | |
| | Group x Time | | | .006 | .007 | .769 | .442 | .884 |
| P-HLEs | | .010 | 12.542 | | | | | |
| | Intercept | | | 17.373 | .176 | 98.830 | .000 | |
| | Group | | | .583 | .251 | 2.329 | .020* | |
| | Time | | | .009 | .005 | 1.777 | .076 | |
| | Group x Time | | | .003 | .007 | .373 | .709 | .709 |

Group refers to low- and high-scoring individuals on the factor Personified HLEs, i.e., participants scoring below or above the median. Corresponding state maps can be found in Table 2; *p < .05; **p < .01 reported p-values are uncorrected.
